# Near-chromosome level genome assembly of the fruit pest *Drosophila suzukii* using long-read sequencing

**DOI:** 10.1101/2020.01.02.892844

**Authors:** Mathilde Paris, Roxane Boyer, Rita Jaenichen, Jochen Wolf, Marianthi Karageorgi, Jack Green, Mathilde Cagnon, Hugues Parinello, Arnaud Estoup, Mathieu Gautier, Nicolas Gompel, Benjamin Prud’homme

## Abstract

Over the past decade, the spotted wing Drosophila*, Drosophila suzukii*, has invaded Europe and America and has become a major agricultural pest in these areas, thereby prompting intense research activities to better understand its biology. Two draft genome assemblies based on short-read sequencing were released in 2013 for this species. Although valuable, these resources contain pervasive assembly errors and are highly fragmented, two features limiting their values. Our purpose here was to improve the assembly of the *D. suzukii* genome. For this, we generated PacBio long-read sequencing data at 160X sequence coverage and assembled a novel, contiguous *D. suzukii* genome. We obtained a high-quality assembly of 270 Mb (with 546 contigs, a N50 of 2.6Mb, a L50 of 15, and a BUSCO score of 95%) that we called WT3-2.0. We found that despite 16 rounds of full-sib crossings the *D. suzukii* strain that we sequenced has maintained high levels of polymorphism in some regions of its genome (ca. 19Mb). As a consequence, the quality of the assembly of these regions was reduced. We explored possible origins of this high residual diversity, including the presence of structural variants and a possible heterogeneous admixture pattern of North American and Asian ancestry. Overall, our WT3-2.0 assembly provides a higher quality genomic resource compared to the previous one in terms of general assembly statistics, sequence quality and gene annotation. This new *D. suzukii* genome assembly is therefore an improved resource for high-throughput sequencing approaches, as well as manipulative genetic technologies to study *D. suzukii*.

## INTRODUCTION

*Drosophila suzukii* (Matsumura, 1931), the spotted wing Drosophila (Diptera: Drosophilidae), is an invasive fruit fly species originating from eastern Asia that has spread since 2008 in major parts of America and Europe. This species is still expanding its distribution (Cini, Ioriratti, & Anfora, 2012; Deprá, Poppe, Schmitz, De Toni, & Valente, 2014) and is classified as a major pest on a variety of berries and stone fruit crops (Asplen et al., 2015). Its behavior and phenotypic traits are now the subject of intense scrutiny both in the lab and in the field (reviewed in (Olazcuaga, Rode, et al., 2019)).

Understanding the biology and the population dynamics of *D. suzukii* benefits from the production and mining of genomic and transcriptomic data, as well as manipulative genetic technologies including functional transgenesis and genome editing (Kalajdzic & Schetelig, 2017; Karageorgi et al., 2017; J. Li & Handler, 2017). Yet, the efficacy of these approaches relies critically on high-quality genomic resources. Currently, two *D. suzukii* genome assemblies, obtained from two different strains, have been generated based on short-read sequencing technologies (Chiu et al., 2013; Ometto et al., 2013). The utility of these valuable genomic resources is limited by the extensive and inescapable assembly errors, as well as the high fragmentation rates that characterize short-read sequencing genome assemblies.

While short read sequencing has dramatically contributed to the vast repertoire of genomes available nowadays, it has the unsolvable issue that ~100 bp-long reads cannot resolve genomic structures with low-complexity or polymorphic regions, and produce a large number of relatively short contigs (e.g. (Chiu et al., 2013)). The advent of long-read sequencing technology (e.g. nanopore, PacBio) that produces reads that are several dozens of kilobases (kb) long on average has proven an efficient tool to circumvent those limitations and allows to assemble much longer contigs at least for small to medium-sized genomes (Gautier et al., 2018; Gordon et al., 2016; Jiao et al., 2017; Korlach et al., 2017; Matthews et al., 2018; Weissensteiner et al., 2017).

Genome assemblies using long read sequencing have been generated for at least 18 Drosophila species (Allen, Delaney, Kopp, & Chenoweth, n.d.; Bracewell, Chatla, Nalley, & Bachtrog, 2019; Mahajan, Wei, Nalley, Gibilisco, & Bachtrog, 2018; Miller, Staber, Zeitlinger, & Hawley, 2018).Somewhat surprisingly given its economic importance, *D. suzukii* is missing from this list. In this article, we report the genome assembly of an inbred *D. suzukii* strain using the long-read Pacific Biosciences (PacBio) technology. The assembly compares favorably to previous ones in terms of general assembly statistics, detailed sequence quality and gene annotation, although some parts, mostly located on regions homologous to the 3R *D. melanogaster* chromosome arm, remained fragmented and displayed high residual genetic diversity in the strain. The improved assembly allowed us to further explore some specific aspects of the *D. suzukii* genome including repeats content, structural variants, sequence diversity and population genetic origins.

## MATERIALS AND METHODS

### Whole-genome long-read sequencing of the WT3-2.0 *D. suzukii* strain

The WT3-2.0 *D. suzukii* individuals used to produce our genome assembly derived, after six additional generations of full-sib crossing, from the WT3 isofemale strain (here named WT3-1.0) that was previously established from a female sampled in Watsonville (USA) and sequenced by (Chiu et al., 2013). The WT3-2.0 strain hence went through a total of at least 16 rounds of full-sib crossing.

#### Genomic DNA extraction

High-molecular weight DNA was extracted from 40 adult D. suzukiii females (WT3-2.0) using the Blood & Cell culture DNA midi kit (Qiagen). The quality and concentration of the DNA was assessed using a 0.5 % agarose gel (run for >8 h at 25 V) and a Nanodrop spectrophotometer (ThermoFisherScientific). PacBio libraries were generated using the SMRTbell™ Template Prep Kit 1.0 according to manufacturer’s instructions. In brief, 10 µg of genomic DNA per library (estimated by Qubit assay) was sheared into 20 kb fragments using the Megaruptor system, followed by an exo VII treatment, DNA damage repair and end-repair before ligation of hair-pin adaptors to generate a SMRTbell™ library for circular consensus sequencing. The library was then subjected to exo treatment and PB AMPure bead wash procedures for clean-up before it was size selected with the BluePippin system (SAGE) with a cut-off value of 9000 bp. In total 48 units of SMRTcell™ with library was sequenced on the PacBio Sequel instrument using the Sequel 2.0 polymerase and 600 minute movie time. The raw data were then imported into the SMRT Analysis software suite (v2.3.0) where subreads shorter than 500 bp and a polymerase read quality below 75 were filtered out.

### Genome assembly based on PacBio long reads

We generated two separate assemblies using two approaches: Falcon (https://github.com/PacificBiosciences/FALCON) using the parameters detailed in the Supplementary Text S1 and Canu 1.3 (Koren et al., 2017) with the default options (except *-minReadLength=7000-stopOnReadQuality=0-minOverlapLength=1000*)

The resulting Falcon assembly, hereafter called *dsu_f*, was 281Mb long while the Canu assembly, hereafter called *dsu_c*, was 267Mb long. We noticed that each assembly lacked different parts of the genome. For instance the gene Abd-B was absent from *dsu_c* and the gene Or7a was absent from *dsu_f*; see Figure S1 for more exhaustive BUSCO gene content statistics (Simão, Waterhouse, Ioannidis, Kriventseva, & Zdobnov, 2015). We therefore decided to follow a hybrid strategy (similarly to (Chakraborty, Baldwin-Brown, Long, & Emerson, 2016) and merged these two assemblies. To that end we proceeded in three successive merging steps (following recommendations provided by Mahul Chakraborty’s) using: (i) the *nucmer* (with options *-l 100*) and *delta-filter* (with options *-i 95-r-q*) programs from the MUMmer v3.23 package (Kurtz et al., 2004) to perform alignment of assemblies on a whole genome scale, and (ii) the *Quickmerge* program (Chakraborty et al., 2016) (with options *-hco 5.0-c 1.5-l 660000-ml 10000*) to merge assemblies based on their resulting alignment. In the first step, we aligned *dsu_c* (taken as reference) and *dsu_f* (taken as query) and obtained the *dsu_fc* merged assembly. In the second step, we aligned *dsu_fc* (taken as reference) and *dsu_c* (taken as query) and obtained the *dsu_fc2* merged assembly. In the third and last step, we aligned *dsu_fc2* (taken as reference) and *dsu_f* (taken as query) and obtained the *dsu_fc2f* merged assembly. We further added a polishing step to account for the high error rate in PacBio reads (above 10% (Korlach, 2013)). This polishing step can be performed after the merging (Solares, G3 2018) using PacBio reads if they are abundant enough (Chakraborty et al., 2016; Chin et al., 2013). We mapped back a subset of our PacBio raw data (to obtain 80X coverage) to the *dsu_fc2f* assembly using *pbalign* and corrected the assembly using *quiver* with default parameters (both programs obtained from the SMRT Portal 2.3; http://www.pacbiodevnet.com). The *dsu_fc2f_p* resulting assembly was ~286Mb long and contained 669 contigs.

We finally sought to remove both exogeneous sequences (e.g., bacterial contaminant) and duplicated sequences resulting from the poor handling of diploidy by assemblers (although *Falcon* produces a partially diploid genome). We first used BUSCO v2 with the bacterial database (bacteria_odb9 containing 148 genes in 3663 species) to identify contigs containing bacterial genes. Twenty-two contigs were removed from the assembly, most of them aligning onto the *Acetobacter pasteurianus* genome. Also, we manually retrieved five additional contigs mapping to *Lactobacillus* genome leading to a total of 27 bacterial contigs discarded (corresponding to ca. 3Mb). We then used BUSCO v2 with the *Diptera* database (diptera_odb9 containing 3295 genes in 25 species) to identify putative duplicated contigs and flagged the shortest one as redundant. To avoid removing valid contigs, we recovered the contigs that contained at least 10 predicted genes. To this end, we mapped possible ORFs (longer than 200bp) using the NCBI tool ORFinder v0.4.0 (https://www.ncbi.nlm.nih.gov/orffinder/) that was run with default options. The identified ORFs were then aligned onto the assembly without the redundant contigs using BLAST (Altschul, Gish, Miller, Myers, & Lipman, 1990), considering as significant hits with e-value<10^−4^ and >80% of identity. Using this procedure, we removed 69 contigs that had fewer than ten unique ORFs considering that they were likely alternative sequences and we split four contigs in two because the unique ORFs were all located at a contig end with the rest of the contig appearing duplicated. Also, the annotation of structural variants (see below) lead us to remove 27 additional contigs, flagged as translocations but that further scrutiny made us consider as alternative haplotypes. In total 96 redundant contigs plus 4 partial contigs (covering ca. 16 Mb) were removed from the main assembly and were added to the file already containing 457 alternative haplotyes assembled by Falcon, covering approximately 26 Mb and assembled together with *dsu_f*. In total, this resulted in an assembly of alternative haplotypes of ca. 42 Mb in total.

Only 16 Mb out of the 42 Mb of alternative sequences were assigned to a contig from the main assembly using BUSCO. In addition, this assignation did not provide precise positioning on the main assembly. We therefore decided to precisely map all alternative sequences to our main *D. suzukii* assembly using the methodology described in the section “Whole genome alignment with other assemblies” below. Aligned regions that varied in size more than two folds between the alternative sequence and the main assembly were filtered out. Following this procedure, we were able to assign 91% of the contigs to the main assembly, covering 97.5% of the alternative sequences (41 Mb / 42 Mb).

The final assembly, hereafter called WT3-2.0 consisted of 546 contigs for an overall size of 268 Mb. The contiguity of the assembly was measured using Quast 4.1 (Gurevich, Saveliev, Vyahhi, & Tesler, 2013) (run with default options) and its completeness was evaluated against the *Diptera* gene set with BUSCO v2 run with option *-c 40*.

### Assessment of local assembly errors

We used the following procedure to identify local genome assembly errors in the form of short (ca. 1kb long) sequences duplicated in tandem. We used the genome assemblies of *D. melanogaster* dm6 (Genbank reference GCA_000001215.4), *D. biarmipes* Dbia_1.0 (GCA_000233415.1), the previous *D. suzukii* WT3-1.0 assembly (GCA_000472105.1) and our assembly WT3-2.0. We selected exons of an annotation that were within 5 kb of each other but did not overlap. We then blasted them against each other and selected hits that aligned on more than 50% of the shortest among the pair with an e-value below 10^−10^. For each couple of retained exons, a Needleman/Wunsch global alignment was made using the nw.align 0.3.1 python package (https://pypi.python.org/pypi/nwalign/) and a score was calculated with a NUC.4.4 matrix downloaded from the ncbi website. The score was normalized by the length of the sequence alignment and ranged from −2 (lowest similarity) to 5 (identical sequences).

### Identification of autosomal and X-linked contigs using a femal-to-male read mapping coverage ratio

To assign contigs of the new WT3-2.0 assembly to either sex chromosomes or autosomes, we compared the sequencing coverage from whole genome short-read sequence data obtained forone female and one male individual (Bidon, Schreck, Hailer, Nilsson, & Janke, 2015; Gautier et al., 2018). Two DNA paired-end libraries with insert size of ca. 350 bp were prepared using the Illumina TruSeq Nano DNA Library Preparation Kit following manufacturer protocols on DNA extracted using the Genomic-tip 500/G kit (QIAGEN) for one female (*mtp_f19*) and one male (*mtp_m19*) sampled in Montpellier (France). Each individual library was further paired-end sequenced on the HiSeq 2500 (Illumina, Inc.) with insert size of 125 bp. Base calling was performed with the RTA software (Illumina Inc.). The raw paired-end reads, available at the SRA repository under the SRR10260311 (for *mtp_f19*) and SRR10260312 (for *mtp_m19*) accessions, were then filtered using fastp 0.19.4 (Chen, Zhou, Chen, & Gu, 2018) run with default options to remove contaminant adapter sequences and eliminate poor quality bases (i.e., with a Phred-quality score <15). Read pairs with either one read with a proportion of low-quality bases over 40% or containing more than five N bases for either of the pairs were removed. After filtering, a total of 78,629,384 (9.379131 Gb with Q>20) and 52,311,302 (6.342157 Gb with Q>20) reads remained available for *mtp_f* and *mtp_m* respectively with an estimated duplication rate of 0.918% and 0.492%, respectively. Filtered reads were then mapped onto the WT3-2.0 assembly using default options of the MEM program from the BWA 0.7.17 software (H. Li & Durbin, 2009; H. Li et al., 2009; Heng Li, 2013). Read alignments with a mapping quality Phred-score < 20 or PCR duplicates were removed using the view (option -q 20) and markdu**p** programs from the SAMtools 1.9 software (Heng Li, 2013), respectively. The resulting total number of mapped reads for *mtp_f19* and *mtp_m19* was 42,304,522 and 33,301,631 reads with a proportion of properly paired reads of 96.6% and 98.0% respectively.

Sequence coverage at each contig position for each individual sequence was then computed jointly using the default options of the depth program from SAMtools 1.9. To limit redundancy, only one count every 100 successive positions was retained for further analysis and highly covered positions (>99.9^th^ percentile of individual coverage) were discarded. The overall estimated median coverage was 18 and 21 for *mtp_f19* and *mtp_m19*, respectively.

To identify autosomal and X-linked contigs, we used the ratio *ρ* of the relative (median) read coverage of contigs between *mtp_f19* and *mtp_m19* (weighted by their corresponding overall genome coverage). The ratio *ρ* is expected to equal 1 for autosomal contigs and 2 for X-linked contigs (Bidon et al., 2015; Gautier et al., 2018). Note that the inclusion of X-linked positions in the overall estimated male genome coverage to compute the weights in the estimation of *ρ* result in a downward bias (the higher the actual length of the X-chromosome, the higher the bias). As a matter of expedience, 226 contigs (out of 546) with a coverage lower than 5X (resp. 2X) in *mtp_m19* (resp. *mtp_f19*) or with less than 100 analyzed positions (i.e., <10 kb) were discarded from further analyses. Conversely, four additional contigs (namely #234, #373, #668 and #638 of length 132 kb, 51 kb, 13 kb and 21 kb respectively) were discarded because they showed outlying coverages (i.e., > Q3 + 1.5(Q3-Q1), where Q1 and Q3 represents respectively the 25% and 75% quantiles of the observed contig coverage distribution) in either *mtp_f19* or *mtp_m19*. The cumulated length of the 316 remaining contigs was 256.1 Mb. Only 11.9 Mb of the WT3-2.0 assembly were hence discarded. We then fitted a Gaussian mixture model to the estimated *ρ* distribution of these 316 contigs, with two classes of unknown means and the same unknown variance. The latter parameters were estimated using the Expectation-Maximization algorithm implemented in the mixtools R package (Benaglia, Chauveau, Hunter, & Young, 2009). As expected, the estimated mean of the two classes µ_1_=0.93 and µ_2_=1.90 were slightly lower than that expected for autosomal and X-linked sequences. Our statistical treatment allowed the classification of 296 contigs (223.5 Mb) as autosomal and 14 contigs (31.9 Mb in total) as X-linked with a high confidence (p-value < 0.01), only *ca.* 14.5 Mb being left unassigned.

### Genome annotation for repetitive elements, structural variants and coding sequences

The repertoires of repetitive elements was assessed for the following xspecies and genome assemblies: *D. melanogaster* dm6 (Genbank reference GCA_000001215.4), *D. suzukii* WT3-2.0, *D. biarmipes* Dbia_1.0 (GCA_000233415.1) and *D. takahashi* Dtak_2.0 (GCA_000224235.2). We used the following procedure for each species separately: initial sets of repetitive elements were obtained using RepeatMasker open-4.0.6 (Smit et al., 2013-2015) with default parameters and the large Drosophila repertoire of all classes of repetitive elements from the Repbase database (Baena-López, Baonza, & García-Bellido, 2005; Bao, Kojima, & Kohany, 2015). The number of bases covered by each type of repetitive element was then computed on the repertoire. The set of complete elements was obtained using the output of the programs RepeatMasker and OneCodeToFindThemAll (Bailly-Bechet, Haudry, & Lerat, 2014), from which the number of bases covered by each type of repetitive elements was extracted.

To detect structural variants, we aligned our filtered PacBio reads against our WT3-2.0 assembly using NGMLR v0.2.6 (Sedlazeck et al., 2018) with the parameter “-i 0.8”. We then used Sniffles v1.0.7 (Sedlazeck et al., 2018) to detect structural variants with the parameter “-s 20 –l 500”. We detected 53 translocations, 59 insertions, 219 deletions 9 inversions and 61 duplications. During this process, 27 contigs identified as translocations rather corresponded to alternative sequences upon manual inspection and were hence removed from the main assembly (see section “Genome assembly based on PacBio long reads”).

To annotate protein coding genes, we used sequence-based gene prediction as well as cDNA evidence. RNA was extracted from antennae, ovipositors, proboscis + maxillary palps and tarsi of WT3_1.0 adults females and from pupal ovipositors (collected at 6h, 24h and 48h after puparium formation) using Trizol (Invitrogen) according to manufacturer’s instructions. In total, 8 libraries were prepared using the Truseq stranded kit (Illumina) according to the manufacturer’s instructions, and were sequenced on a Hiseq2500.

We used Maker v2.31.8 (Holt et al., 2011) to annotate the genome. SNAP (Korf, 2004) and AUGUSTUS v3.2.2 (Keller et al, 2011 Bioinformatics) with the parameter “augustus_species=fly » were used for *ab initio* predictions. cDNA evidence was provided from Trinity v2.3.2 (Haas et al., 2011) and hisat v2.0.4 (Kim, Langmead, & Salzberg, 2015) plus stringtie v 1.2.4 runs (Pertea et al., 2015) on the RNAseq data on pupae and the different tissues of female adults. We used the *D. melanogaster* proteome as protein homology evidence. The repeatmasker parameter was set to “Drosophila” and a general set of 24,916 transposable element proteins was provided. SNAP was trained with two initial runs: the first run used the homology and cDNA evidence (est2genome and protein2genome were set to 1) and the second run used the SNAP HMM file produced after the first maker run. The final maker run combined all the evidence, trained SNAP parameters as well as AUGUSTUS. In order to correct the flawed tendency of assemblers to fuse two neighboring genes together (Grabherr et al., 2011), we added the following step. A *D. suzukii* gene was called a “false chimeric” if it could be cut into two parts that each aligned by BLAST to two neighboring genes in *D. melanogaster*. Based on this criterion, 1052 genes were identified as chimers and split in two at the position that mimicked the gene limits in *D. melanogaster*. This method is conservative as it implies that gene structure is conserved between species. This assumption is reasonable owing to the evolution of genes and genomes on the Drosophila phylogeny and was validated by a manual review of modified genes (Consortium, 2007).

### Whole genome alignment with other assemblies

The genome sequences of our *D. suzukii* assembly WT3-2.0, the previous *D. suzukii* WT3-1.0 assembly (GCA_000472105.1), the *D. biarmipes* Dbia_1.0 assembly (GCA_000233415.1) and the *D. melanogaster* dm6 assembly (Genbank reference GCA_000001215.4) were aligned as previously described (Paris et al., 2013). Briefly, we followed the general guideline described in (Dewey, 2007): we used a large-scale orthology mapping created by Mercator (Dewey, 2006) with the option to identify syntenic regions of the genomes. Each region was then aligned with Pecan (Paten, Herrero, Beal, Fitzgerald, & Birney, 2008) with default parameters.

To visualize synteny blocks between the 20 longest contigs of the *D. suzukii* assembly and *D. melanogaster* chromosomes, we proceeded as follow. We first ran BLASTP between the protein anchors of *D. melanogaster* and *D. suzukii* produced during genome alignment with the parameters “-e 1e-10-b 1-v 1-m 8”. We then ran MCScanX (Wang et al., 2012) on the BLASTP output using default parameters. Synteny plots were obtained using VGSC (Xu et al., 2016) on the MCScanX output. The results were fully consistent between this method and the genome alignment described above.

### Estimating nucleotide diversity in the original WT3 strain, the WT3-2.0 strain and their populations of origins

We relied on Pool-seq short-read whole-genome shotgun sequencing data (WGS) to estimate nucleotide diversity in the original WT3 strain (Chiu et al., 2013) (here referred to as WT3-1.0), the newly generated WT3-2.0 strain and three wild populations sampled in Watsonville - USA (US-Wat), Hawaii – USA (US-Haw); and Ningbo - China (CN-Nin). These choices were motivated by the fact that the female initially used to establish the WT3 strain originates from the Watsonville population (Chiu et al., 2013) and that the later population has recently been shown to be of admixed origin between Hawaii and Eastern China (Ningbo) populations (Fraimout et al., 2017). For the original WT3 strain (WT3-1.0), WGS data of a pool of tens of females (Chiu, personal communication) used to build the previous genome assembly by (Chiu et al., 2013) was downloaded from the SRA under the accession SRR942805. These Pool-seq data were obtained after sequencing of a DNA paired-end (PE) library with insert size of 250 bp on a HiSeq2000 (Illumina, Inc.) sequencers at approximately 40X coverage (see Table S1 in (Chiu et al., 2013)). For the WT3-2.0 strain, a pool of 26 individual genomes (13 males and 13 females) was sequenced on a HiSeqX sequencer (Macrogen Inc., Seoul, South Korea) targeting a coverage >30X. The raw paired-end sequences (2×150) were made available from the SRA repository under the SRR10260310 accession. Finally, for the US-Wat, the US-Haw and the CN-Nin populations, we relied on the Pool-seq data recently produced by (Olazcuaga, Loiseau, et al., 2019) from samples of 50 individuals (including 4, 25 and 36 females, respectively) and available from the SRA under the SRR10260026, SRR10260031 and SRR10260027 accessions respectively. The three data sets consisted of 2×125 bp PE sequences obtained from a HiSeq 2500 sequencer. Processing and mapping of reads was carried out as described above for the *mtp_f19* and *mtp_m19* individual WGS. The resulting overall mean coverages were 29.8X, 34.5X, 52.7X, 51.7X and 71.8X for WT3-1.0, WT3-2.0, US-Wat, CN-Nin and US-Haw, respectively.

Nucleotide diversity (θ=4Neµ) was then estimated for non-overlapping 10 kb windows across the genome using the extension of the Watterson estimator (Watterson, 1975) for Pool-Seq data developed by Ferretti et al. (2013) and implemented in the npstats software. Only positions covered by at least four reads and less than 250 reads with a min quality >20 were considered in the computations (-mincov 4-maxcov 250-minqual 20 options) and windows with less than 9,000 remaining positions were discarded. As a matter of expedience, the haploid pool sample size was set to 50 individuals for the WT3-1.0 strain. We found, however, that alternative values of 10, 20 or 100 individuals resulted in highly consistent estimates.

### Estimating the local ancestry composition of the assembly using an HMM painting model

Because of the admixed origin of the Watsonville population from which the WT3-1.0 and WT3-2.0 strains originate, we expected the assembly to be a mosaic of chromosomal segments of ancestral individuals originating from Hawaii (US-Haw) and Eastern China (CN-Nin), its two source populations (Fraimout et al. 2017). To characterize this mosaic, we first called polymorphic sites in the US-Haw and CN-Nin populations. To that end, the US-Haw and CN-Nin Pool-seq BAM files (see above) were processed using the mpileup program from SAMtools 1.9 with default options and *-d 5000* and *-q 20*. Variant calling was then performed on the resulting mpileup file using VarScan mpileup2cns v2.3.4 (Koboldt et al., 2012) with options *–min-coverage 50*; *–min-avg-qual 20* and *–min-var-freq 0.001 –variants –output-vcf*. The resulting VCF file was processed with the *vcf2pooldata* function from the R package poolfstats v1.1 (Hivert, Leblois, Petit, Gautier, & Vitalis, 2018) retaining only bi-allelic SNPs covered by >20 and <250 reads in each of the two sample. For each SNP, we then estimated the frequency of the reference allele (i.e., the one of the assembly) in each population using a Laplace estimator (see Supplementary Text S2). We only retained SNPs displaying an absolute difference in the reference allele frequencies above 0.2 between the two US-Haw and CN-Nin samples (i.e. the most ancestry informative SNPs). This resulted in a total of 2,643,102 autosomal and 540,277 X-linked SNPs.

We further developed a one-order Hidden Markov Model (HMM) to model the assembly as a mosaic of chromosomal segments from either Chinese (“C”) or Hawaiian (“H”) ancestry. This HMM allowed estimation of the local ancestry origin of each reference allele of the assembly based on its estimated frequencies in the CN-Nin and US-Haw samples used as proxies for the “C” and “H” ancestral populations respectively. The model and the parameter estimation method are detailed in Supplementary Text S2.

## RESULTS

### Available *D. suzukii* genomic resources contain pervasive local assembly errors

The initial genome assembly of an Italian *D. suzukii* strain, inbred for 5 generations, was highly fragmented (e.g. N50 of 4.5kb for the contigs, L50 of 8700 (Ometto et al., 2013)). In parallel, a second strain, WT3, established from a single female collected in Watsonville, CA, U.S.A, and inbred for 10 generations by sib-mating, had been sequenced (Chiu et al., 2013). The latter assembly, hereafter called WT3-1.0, was more contiguous than the assembly by Ometto et al., (2013) as judged by summary statistics (e.g. N50 of ~27 kb for the contigs and ~385 kb for the scaffolds, and L50 of 73 for the scaffolds; Chiu et al. 2013). Nevertheless, we recurrently observed inconsistencies between the WT3-1.0 assembly and Sanger sequencing data we obtained for specific loci amplified using PCR from the *D. suzukii* strain used by Chiu et al. (2013). For instance, BLAST alignments of the *Orco* locus between the WT3-1.0 assembly and the reference *D. melanogaster* assembly dm6 indicated that exons 2 and 3 were repeated twice in the middle of the gene (Figure 1A), a feature not confirmed by Sanger sequencing data. We suspected that the published genome sequence contained a local assembly error corresponding to a ~1kb long region incorrectly repeated twice. To test if such a pattern of locally duplicated exons was widespread in the WT3-1.0 assembly, we aligned with BLAST all the annotated exons against each other, only retaining hits that were within a 5kb window of each other and with a near-perfect score (score > 4.8 on a scale ranging from 1 to 5). For comparison, the same procedure was applied to assemblies of *D. melanogaster* and *D. suzukii*, and of the sister species *D. biarmipes* (Figure 1B). We only found a few locally duplicated exons in the *D. melanogaster* assembly and almost none in the *D. biarmipes* assembly. Conversely, the WT3-1.0 assembly contained thousands of neighboring, nearly identical exons (corresponding to ~1000 transcripts and ~600 genes), i.e. at least 10 times more than in *D. melanogaster* and 50 times more than in *D. biarmipes*. Assuming that the genome of *D. suzukii* contains similar levels of near identical adjacent exons as *D. melanogaster* or *D. biarmipes*, we suspected that the *D. suzukii* WT3-1.0 assembly may contain many local assembly errors. Such errors could be caused, at least partly, by both a high level of residual polymorphism in the sequenced *D. suzukii* strain, and by the limitations of the short-read sequencing technology.

**Figure 1:**
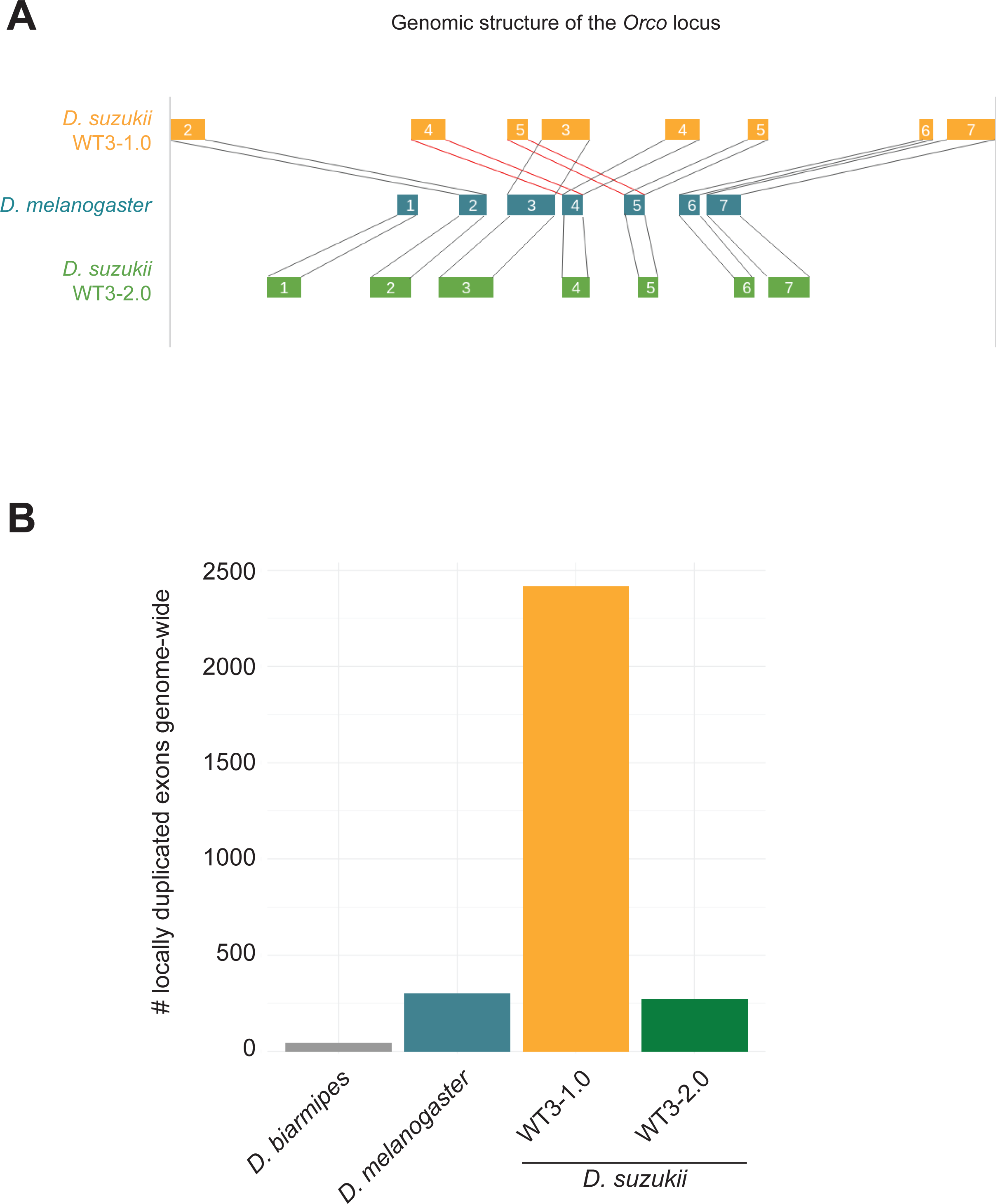
**A.** Genomic structure of the *Orco* gene in the WT3-1.0 genome assembly (Chiu et al., 2013) and in the WT3-2.0 assembly (this article). Genomic structure in *D. melanogaster* is shown for comparison. The locus encompassing exon 1 is missing in the WT3-1.0 assembly. **B**. Number of nearly identical neighboring exons present in *D. suzukii* assemblies WT3-1.0 and WT3-2.0, and in the *D. biarmipes* Dbia_1.0 and *D. melanogaster* dm6 assemblies.

### Long-read sequencing and *de novo* genome assembly

To reduce the genetic diversity of the WT3 strain used by (Chiu et al., 2013) for the WT3-1.0 assembly (Chiu, 2013), we further isogenized flies from this strain by processing full-sib crosses for six generations, resulting in a total of at least 16 generations of inbreeding. We named this new *D. suzukii* strain WT3-2.0 and sequenced genomic DNA extracted from 40 WT3-2.0 females to a coverage of 160x using the single molecule real time sequencing on the Pacific Biosciences technology platform.

We followed a customized approach for the assembly step (see Materials and Methods for details), paying special attention to both bacterial contamination and the putative presence of different haplotypes of the same locus assembled as separate sequences. The resulting assembly consisted of 546 contigs for an overall size of ~270Mb, whichis closer to the estimated genome size of ~313Mb (Sessegolo, Burlet, & Haudry, 2016) than previous assemblies (cf. 232 Mb in Chiu et al., 2013 and 160 Mb in Ometto et al., 2013). Assembly statistics likewise suggest an improvement (N50 of 2.6Mb, L50 of 15, BUSCO score of 95%), both in terms of continuity and completeness (see Table S1 for assembly statistics and Figure S1 for BUSCO results), which indicates substantial improvements over the WT3-1.0 assembly (Table S1). Importantly, the correct non-duplicated structure of the *Orco* gene was recovered and, more generally, the number of locally duplicated exons in the WT3-2.0 assembly was similar to that observed in the *D. melanogaster* assembly (Figure 1).

For approximately 34Mb of the WT3-2.0 assembly two haplotypes of homologous sequence were inferred (see Materials and Methods) whereas the rest of the assembly was identified as haploid. Distinguishing alternative sequences (or haplotypes) from recent duplicates is notoriously difficult. The coverage at regions that have been assembled as one versus regions with two haplotypes was similar (Figure S2A) providing evidence for the former.

Fifty contigs of the WT3-2.0 assembly could be unambiguously aligned onto the *D. melanogaster* dm6 genome assembly. Those 50 contigs covered ~153Mb (~57% of the assembly, ~82% of the annotated genes) and corresponded to most of the largest contigs enriched for protein coding genes. The small contigs tended to be more covered by repetitive elements that could not be assigned a clear orthologous sequence in *D. melanogaster*. For all but one out of those 50 contigs, over 99% of the aligned section matched a unique *D. melanogaster* chromosome. Even the near-chromosome length contigs almost fully aligned to a unique *D. melanogaster* chromosomal arm (e.g., the 26 Mb-long contig1 aligned to 2L and the 25Mb-long contig2 aligned to 3L; Figure 2). This result suggests that few inter-chromosomal rearrangements occurred since the last common ancestor of *D. melanogaster* and *D. suzukii*. Still, sequence similarity with *D. melanogaster* provides a reliable estimate of the chromosomal origin for most *D. suzukii* contigs (Figure 2).

**Figure 2:**
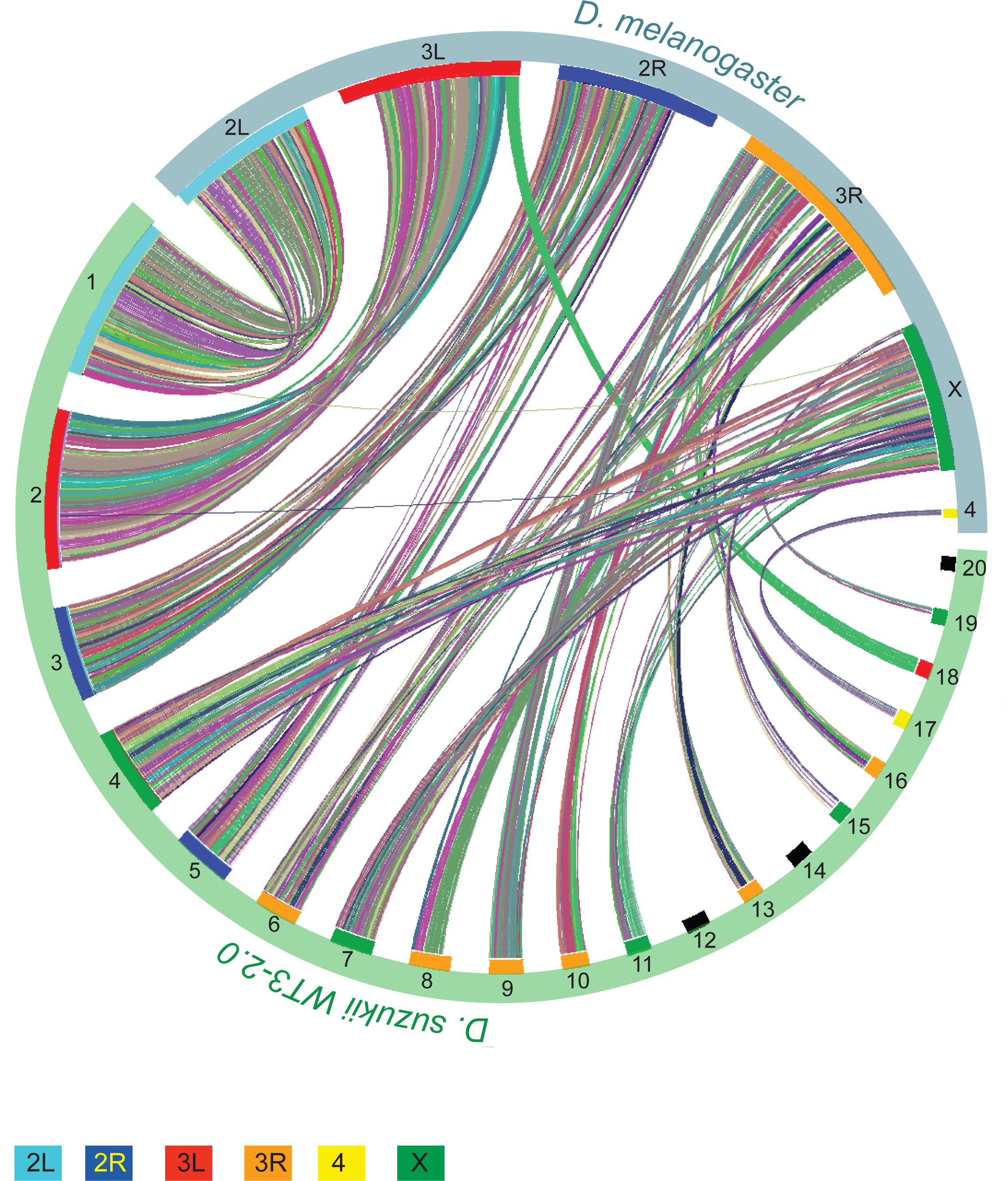
Co-linearity between the 20 longest contigs of the WT3-2.0 assembly and *D. melanogaster* chromosome arms. Colors represent synteny blocks (automatically assigned by VGSC.

Next, we sequenced one male and one female with short-reads at approximately 20X coverage to assign contigs to either autosomes or the X chromosome based on the ratio of male to female read coverage. This confirmed previous contig assignment and lead to the assignment of six additional contigs (totaling 337 kb) to the X-chromosome and 258 additional contigs (totaling 101 Mb) to autosomes. The remaining 234 contigs (amounting to 13 Mb and hence representing less than 5% of the assembly) could not be assigned to autosomes or the X chromosome. The consistency between both assignment methods (sequence synteny with *D. melanogaster* and male over female genomic read coverage) suggested that no large-scale translocations between the X and the autosomes occurred since the split between *D. suzukii* and *D. melanogaster* genomes. Accordingly, nomenclature of *D. suzukii* contigs was based on synteny and contigs were named after the arm to which they align on the *D. melanogaster* genome. Because the WT3-2.0 assembly was obtained from female DNA only, the Y chromosome could not be sequenced and assembled.

### Genome Annotation

We compared the content of various categories of repeated elements in our new *D. suzukii* assembly with that of other Drosophila species, including *D. melanogaster*, *D. biarmipes* and *D. takahashi*. We found that our *D. suzukii* assembly has a particularly large repeatome (~93 Mb corresponding to more than 35% of our 270 Mb-long genome assembly, Figures S3A and S3B), which was twice the size of the repeatome estimated from the previous WT3-1.0 *D. suzukii* assembly. The repeatome in a *D. melanogaster* assembly made from the same PacBio technology (Chakraborty et al., 2018) was about half less, amounting to 45Mb in size and corresponding to 25% of the genome. Likewise, the repeatome was around 21% of the genome in both *D. biarmipes* and *D. takahashi*. Overall, the ~50 Mb inflation of repetitive elements in the ~270 Mb *D. suzukii* genome compared to the ~150 Mb-long *D. melanogaster* genome represented about half of the genome expansion in *D. suzukii*.

We assessed whether this increase in repetitive sequences was coupled with a change in element repertoire. There are different types of repetitive elements such as long satellite DNA, terminal repeat (LTR) retrotransposons, LINE (long interspersed nuclear element)-like retrotransposons, terminal inverted repeat (TIR) DNA-based transposons, and satellite elements (Kaminker et al., 2002). The different types of elements were found in the *D. suzukii* genome in proportions similar to those found in the genome of other *Drosophila* species, to the noticeable exception of *D. melanogaster* (Figures S3C, S3E, S3G, S3I). The detailed repertoire from each class was more variable (Figures S3D, S3F, S3H, S3H).

We finally annotated the genome for coding sequences using three sources of information: (i) *de novo* predictions, (ii) sequence similarity with *D. melanogaster* gene annotations, and (iii) *D. suzukii* RNAseq data that we produced from several embryonic and adult tissues. For compact genomes, gene prediction methods tend to annotate neighboring genes as erroneous chimers (Martin and Wang 2011), an issue we also encountered. To partially fix this problem, we systematically searched and corrected those erroneous fissions and also manually curated about 50 genes of particular interest. The resulting annotation was composed of 18,241 genes, 10,557 of them showing a clear orthology with *D. melanogaster* genes using our genome alignment (Table S2). Out of the 10,557 genes found in both WT3-2.0 and *D. melanogaster*, 8,576 were also found in the annotation of the WT3-1.0 assembly (Chiu et al 2013). The 1,981 missing genes were mostly located in genomic regions that were poorly assembled or absent in the WT3-1.0 assembly.

### Understanding pervasive fragmentation of some parts of the assembly

Although improved, the quality of the WT3-2.0 assembly differed along the genome. In particular, the contigs matching chromosome arm 3R were poorly assembled and were the most fragmented part of our assembly (Figure 2). The longest contig associated with 3R was relatively short (~6.5Mb), contributing to ~17% of the total length of contigs associated with 3R, compared to ~15Mb to 26Mb (i.e., 44 to 93% of total length) for other chromosomes. In addition, the WT3-2.0 assembly contained ~12Mb of regions with high rates of local sequence errors in the form of small indels, mostly located in the same genomic areas as the regions assembled as distinct haplotypes, that is on chromosome 3 (76% on 3R and 18% on 3L; Figure 3).

**Figure 3:**
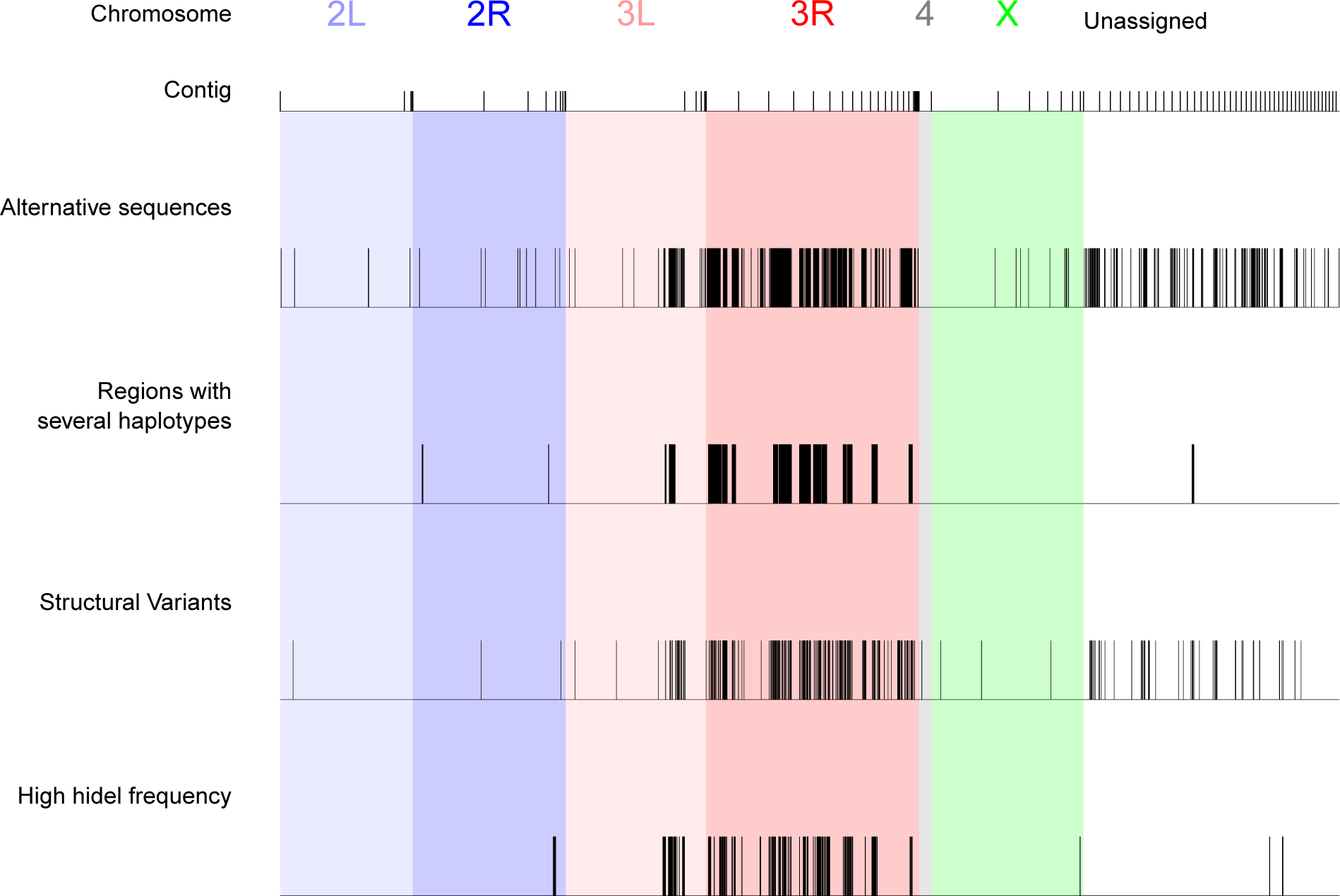
Location at the contig level of various genomic features on the WT3-2.0 assembly. Regions assembled as distinct haplotypes, regions of higher nucleotide diversity, regions with structural variants, regions with higher one-nucleotide assembly errors (in the form of indels) are highlighted. The criteria for defining the boundaries of “high indel rate” and “high nucleotide diversity” regions were as follows: a region is initialized if the rate is above 0.005 on at least 5 consecutive windows of 10 kb; the region is closed if the rate drops below 0.001 on at least 5 consecutive 10 kb windows. Using this rule, 64 regions with high indel rates (median length 140 kb) and 27 regions with high nucleotide diversity (median length 420 kb) were identified. Contigs are ordered according to their chromosomal assignment. Only the longest 100 contigs are shown, representing 79% of the total length of the assembly.

About 34Mb of the WT3-2.0 genome was assembled as two distinct haplotypes (see above), suggesting that residual genetic diversity was maintained in some regions of the *D. suzukii* strain we sequenced, despite a total of 16 rounds of full-sib crossing. We tested this hypothesis by characterizing the patterns of nucleotide diversity estimated from the sequencing of pools (Pool-seq) of 26 individuals from the WT3-2.0 strain. We confirmed that the regions had a higher nucleotide diversity compared to other genomic regions (Figure S2B). This was also true when focusing on chromosome 3R (Figure S2C).

(Fraimout et al., 2017) demonstrated that the population of Watsonville (USA), from which the inbred strain WT3 was derived, originated from an admixture between the native population from Ningbo (China) and the invasive population from Hawaii (USA). This hybrid origin may have resulted in heterogeneity in the distribution of genetic diversity in the genome, maintained in the WT3-2.0 assembly despite strong inbreeding. To test this hypothesis, we first used Pool-seq data to compare the patterns of nucleotide diversity in (i) the WT3-1.0 strain (data from (Chiu et al., 2013)), (ii) the WT3-2.0 strain (see above), (iii) a population sample of the Watsonville area (US-Wat) from which the WT3-1.0 and WT3-2.0 strains originate, and (iv) the aforementioned two source populations of the admixed population from Watsonville (US-Haw and US-Nin). As expected, the genome-wide autosomal nucleotide diversity was maximal in the native Chinese population Ningbo (θ=29.2×10^−3^), lower in the introduced invasive population from Hawaii (θ=12.7×10^−3^), and intermediate in the admixed population of Watsonville (θ=18.2×10^−3^) (Figure 4A). The diversity was lower on the X-linked regions compared to the autosomal regions in all these three populations but the ranking was the same with θ=19.2×10^−3^, θ=6.61×10^−3^ and θ=9.94×10^−3^ for the Ningbo, Hawaii and Watsonville populations, respectively. As expected, the nucleotide diversity on both the autosomes and the X was strongly reduced in the WT3-1.0 strain (θ=5.07×10^−3^ and θ=0.340×10^−3^ respectively), as a result of ten generations of full-sib crossing, and even further reduced (for autosomes) in the WT3-2.0 strain (θ=4.48×10^−3^ and θ=0.420×10^−3^, respectively), which underwent six additional generations of full-sib crossing (Figure 4A). Interestingly, at a chromosomal genomic scale, the distribution of nucleotide diversity was heterogenous, with contigs mapping to chromosome arms 3L and 3R showing substantial residual diversity in the WT3-1.0 strain with estimated θ=4.88×10^−3^ and θ=5.50×10^−3^, respectively, while θ < 0.50×10^−3^ in contigs assigned to other chromosome arms. In the WT3-2.0 strain, the nucleotide diversity dropped to θ=0.39×10^−3^ in 3L, but 3R conserved almost the same levels of diversity as in the WT3-1.0 strain (θ=4.07×10^−3^). These results therefore support our hypothesis that residual genetic diversity is maintained in *D. suzukii* strain WT3-2.0, notably on chromosome arm 3R. Accordingly, 89% of the diploid regions that could be assigned to a *D. melanogaster* chromosome mapped to 3R.

**Figure 4:**
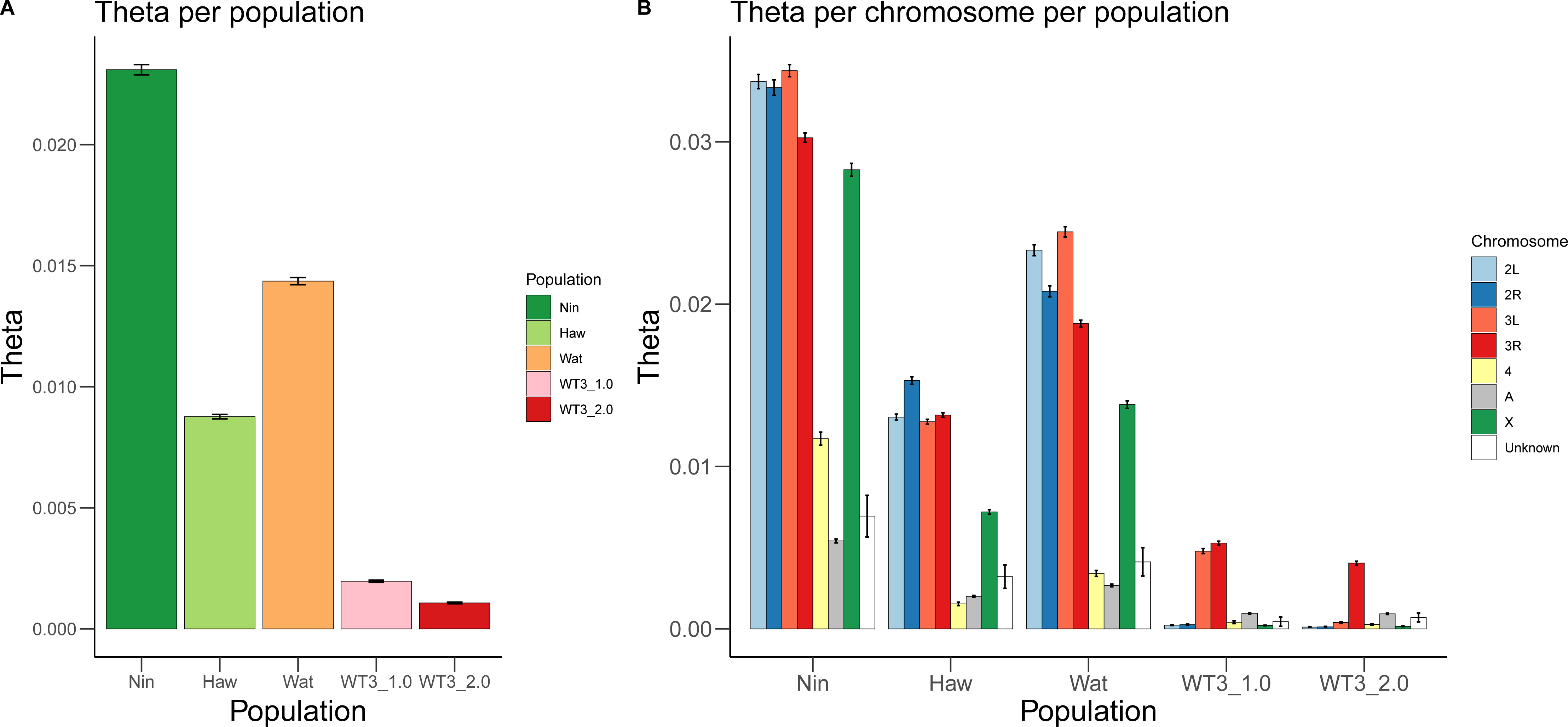
Comparison of nucleotide diversity among *D. suzukii* strains and populations, and among chromosomes. Nucleotide diversity was estimated from pools of individuals for a Chinese population (Nin), a Hawaiian population (Haw), the Watsonville population (Wat), the WT3-1.0 strain (Chiu et al., 2013) and the WT3-2.0 strain (this study). Values of nucleotide diversity parameter Theta (θ) were estimated over 10 kb windows (**A**) genome-wide or (**B**) per chromosome. “2L”, “2R”, “3L”, “3R”, “4”, “X”: assigned *D. melanogaster* chromosome for each WT3-2.0 contigs. “A”: autosomal contigs with no clear corresponding *D. melanogaster* chromosome. “Unknown”: contigs for which chromosomal features (i.e., assignment to autosomal or X chromosomes and to *D. melanogaster* chromosome arm) remain unknown. Error bars correspond to S.E.M.

Next, we tested whether the elevated sequence diversity on chromosomal arm 3R in the WT3 and WT3-2.0 strains could be explained by a heterogeneous pattern of local ancestry origin by characterizing the relative contributions of the Chinese and Hawaiian ancestries to the genome assembly, both at a global and at a chromosomal genomic scale. More specifically, we developed a Hidden Markov Model to determine the genetic origin of the assembled genome at each position using the Pool-seq data produced for the two populations Ningbo and Hawaii mentioned above (see Supplementary Text S2). Overall, the mean fraction α of the assembled genome with a Ningbo origin was found equal to 0.784 (SD = 1.86×10^−3^) for autosomal regions and 0.763 (SD = 8.52×10^−3^) for X-linked regions. We found no significant differences in α values among the different chromosomes (Figure S4). Remarkably, these α values were close to those found in a previous study based on microsatellite markers for the wild source population from Watsonville (i.e. α = 0.759, 90% credibility interval [0.659; 0.854]) (Fraimout et al., 2017). The relative proportions of Chinese and Hawaiian ancestry characterizing the source population from Watsonville have been thus globally preserved in the WT3-2.0 strain. The heterozygous regions, which are mostly concentrated on chromosome arm 3R, only showed a very mild difference in their inferred Hawaiian/Ningbo origin compared to the other regions of the genome (Figure S2D). This means that the elevated nucleotide diversity on 3R (Figure S2B and 2C) cannot be explained by a peculiar pattern of admixture, where for instance a Hawaiian ancestry would have been preferentially retained in these genomic regions.

Finally, we tested whether an excess of structural variants on 3R could contribute to the higher levels of residual polymorphism in the contigs mapping to this particular chromosome arm. Structural variants (SV) are an important source of large-scale polymorphism (e.g. (Joron et al., 2011) but are difficult to detect using short sequencing reads and have been essentially studied by comparing populations that carry different fixed variants (e.g. (Lamichhaney et al., 2016; Tusso et al., 2019), Weissensteiner in prep). To study SVs in our WT3-2.0 strain, we took advantage of PacBio long reads, which give unprecedented access to such information. We detected a total of 369 SVs, mostly Copy Number Variations (338 CNVs including 219 deletions, 59 insertions and 60 duplications), 23 translocations and 8 inversions. We found that the SVs co-localized well with the highly polymorphic regions (Figure 3). We noted that the inversions were of relatively large size, with two inversions longer than 100 kb, (compared to 21 kb in *D. melanogaster*,(Chakraborty et al., 2018)) and an average size of~40 kb (compared to 21 kb in *D. melanogaster*). We confidently assigned six inversions to a *D. melanogaster* chromosome and found that three of them were located on 3R (including one of the longest). In addition, we could confidently assign a *D. melanogaster* chromosome to both extremities of 10 translocations, out of which five were between contigs of different chromosomes, and five between contigs of 3R. This is probably because 3R is the most fragmented chromosome in our assembly, so both extremities of the translocation event are located on two different contigs that belong to the same chromosome but have not been assembled together. Those translocations were located at the end of contigs and are thus probably not real inter-chromosomal translocations but rather either contiguous contigs on 3R or very large inversions within 3R. Altogether, these results are consistent with the hypothesis that the maintenance of structural variants contributes to the residual sequence polymorphism in some areas of the WT3-2.0 genome assembly.

## DISCUSSION

In this article, we used the PacBio long-read technology to re-sequence, assemble and annotate the genome of *D. suzukii*, an invasive fly species that has caused agricultural damage worldwide. This genome was presumably more difficult to assemble than most other Drosophila genomes because of its larger size and higher content of repetitive elements (Sessegolo et al., 2016). It is hence not surprising that the previous *D. suzukii* genome assemblies based on short-read sequencing technologies (Chiu et al., 2013; Ometto et al., 2013) contained pervasive assembly errors. The long-read assembly presented here constitutes a clear improvement which is in line with other assemblies obtained using the long-read technology which bypasses the limitations of short-read sequencing (e.g. (Bracewell et al., 2019; Chakraborty et al., 2018; Miller et al., 2018)). Besides improving general assembly statistics, we made this assembly as workable as possible, notably for molecular biologists. To this aim, we paid special attention to assembly errors caused by contamination or a poor handling of polymorphism by assembly tools.

We also produced a gene annotation that could be compared with *D. melanogaster* (orthology table given as Table S2). Although our *D. suzukii* assembly corresponds to the second largest Drosophila genome after *D. virilis* (333 Mb; (Gregory & Johnston, 2008)), it has a similar gene content compared to *D. melanogaster*, a feature observed so far for all sequenced Drosophila species; (e.g. (“Evolution of genes and genomes on the Drosophila phylogeny,” 2007)). We found that the *D. suzukii* genome contains a high amount of repetitive sequences, as previously shown using a genome-assembly free approach (Sessegolo et al., 2016). Our results suggest that this expansion of the repeatome is responsible for at least half of the increase in genome size in *D. suzukii* (roughly +100 Mb in our assembly), as compared to the closely-related species *D. melanogaster*.

We found that, although nucleotide diversity was globally strongly reduced in the inbred strain WT3-2.0 (and to a lesser extent in the strain WT3-1.0) compared to the wild source population from Watsonville, the residual diversity was heterogeneously distributed among chromosomes, with the highest levels observed on the chromosome 3R homolog for WT3-2.0 (in 3R and 3L for WT3-1.0). This higher residual polymorphism of chromosomal arm 3R is probably responsible, at least partly, for the reduced assembly quality in this genomic region. Because PacBio reads have a very high error rate (~15%, mostly insertions; (Carneiro et al., 2012)), the assembly algorithms that we used tend to interpret heterozygous SNPs as sequencing errors (i.e., insertions) to be removed. Thus, this results in an “overpolished” assembly that contains small errors in the form of single nucleotide deletions (personal communication from PacBio). In agreement with this, we did detect a higher rate of indels on 3R. As a consequence, special caution should be observed for regions of high polymorphism because they tend to display higher assembly error (in the form of one-nucleotide indels).

The substantial level of residual nucleotide diversity in the WT3-2.0 strain on the chromosome 3R homolog remains puzzling. Large inversions have regularly been shown to maintain polymorphism because they prevent recombination between paired loci (e.g., (Joron et al., 2011), (Thomas et al., 2008)). We detected some inversions on 3R, but they are too small to fully account for the high residual sequence polymorphism on this chromosome. However, large inversions are difficult to identify on 3R because they would likely appear as translocations since both extremities of the inversions would end up on different contigs. In agreement with this, many of the translocations that we detected involved two contigs located on 3R. However, the current state of our assembly does not allow us to provide a clear answer. Assembling chromosome 3R from the genome of the parental populations (i.e., Hawaii and Ningbo), in which such long inversions might be absent or homozygous, may help solving this issue because those would be homozygous for the inversions and thus easier to assemble. Long inversions could also be searched using methods that detect physical linkage between regions of the genome (e.g. Hi-C, (Bracewell et al., 2019; Dudchenko et al., 2017; Harewood et al., 2017)). Our results also suggest that despite the tremendous progress in sequencing technology, the complexity and diversity of genomic structures and sequences, even within an isogenized strain, might make fully chromosome-length assemblies difficult to reach for some regions in some species, and the problem worsens for wild individuals.

From an evolutionary perspective, the maintenance of nucleotide diversity in specific regions of the genome may be linked to balancing selection. Both theoretical and empirical works have recently suggested that enhanced neutral variability at nucleotide sites may be closely linked to sites under long-term balancing selection (i.e. sites for which heterozygous individuals have an advantage over homozygous individuals; (Croze et al., 2017; Gao, Przeworski, & Sella, 2015; Zhao & Charlesworth, 2016)). Balancing selection is indeed a selective process that actively maintains multiple variants of genes at frequencies larger than expected from genetic drift alone. Although not studied so far in *D. suzukii*, experimental evidence in several other Drosophila species (especially in *D. melanogaster*) confirms the maintenance of high level of variability in specific genomic regions through balancing selection (Charlesworth, 2015; Croze et al., 2017). Moreover, it has been reported that the level of variability in both marker loci and quantitative traits in laboratory populations can sometimes decline considerably less rapidly over time than is expected under the standard neutral model (e.g., (Gilligan, Briscoe, & Frankham, 2005) in *D. melanogaster* and (Williams et al., 2016) in cattle). Theoretical studies show that, in small (e.g., laboratory reared) populations, associative overdominance due to strong linkage disequilibrium with sites under balancing selection can explain the maintenance of sequence variability in some genomic regions (e.g. (Zhao & Charlesworth, 2016)). In *D. suzukii*, the lower decline of nucleotide diversity observed on chromosome 3, compared to the rest of the genome, might be due to a higher frequency of sites under long-term balancing selection and/or lower recombination rates increasing linkage disequilibrium with those sites. In any case, potential heterozygous advantages do not seem to be associated with genetic admixture, as we found that the chromosome 3R did not show any atypical admixture pattern in our *D. suzukii* assembly. More analyses are needed to confirm the potential link between high balancing selection and the chromosomal heterogeneity of nucleotidic diversity observed in WT3-2.0 but also in WT3-1.0 and in wild populations of *D. suzukii*.

## Conclusion

Our WT3-2.0 assembly provides a higher quality genomic resource compared to the previous one. It confirms the benefits of long-read sequencing for *de novo* assembly. As a short-term perspective, we anticipate that our near-chromosome level assembly should be amenable to a chromosome-level assembly. In particular, scaffolding methods using Hi-C data will represent one of the most promising routes to this purpose. We believe that our improved *D. suzukii* assembly will provide a solid genomic basis to investigate basic biological questions about *D. suzukii*, using high-throughput sequencing technologies as well as manipulative genetic technologies (McCartney, Mallez, & Gohl, 2019).

## ACKNOWLEDGMENTS

We thank D. Begun for providing us with flies of the WT3 strain, Lasse Bräcker for further inbreeding of the *D. suzukii* WT3 stock, Mahul Chakraborty for useful suggestions on the assembly procedure of PacBio long reads, and Tom Druet for helpful discussions on HMM modeling. MP, MK, JEG and BP acknowledge financial support from the CNRS, the European Research Council under the European Union’s Seventh Framework Programme (FP/2007-2013) / ERC Grant Agreement n° 615789; NG and JW acknowledges funding from the Ludwig-Maximilians University of Munich; AE & MG acknowledge financial support by the National Research Fund ANR (France) through the project ANR-16-CE02-0015-01 (SWING), the Languedoc-Roussillon region (France) through the European Union program FEFER FSE IEJ 2014-2020 (project CPADROL), and the INRA scientific department SPE (AAP-SPE 2016 and 2018). JW acknowledges support of the National Genomics Infrastructure (NGI) / Uppsala Genome Center and UPPMAX for providing assistance in massive parallel sequencing and computational infrastructure. Work performed at NGI / Uppsala Genome Center has been funded by RFI / VR and Science for Life Laboratory, Sweden

## DATA ACCESSIBILITY

The PacBio reads and the RNA-seq reads were submitted to SRA under the Bioproject accession number PRJNA594550. The detailed SRA accession numbers are as follows:

### PacBio data (SRA accession numbers)

- SRR10716756 to SRR10716759, SRR10716769 and SRR10716772 to SRR10716814

### RNA-seq data (SRA accession numbers)

- SRR10716760 for male genital discs, 6h after puparium formation

- SRR10716761 for female genital discs, 6h after puparium formation

- SRR10716762 for male genital discs, 48h after puparium formation

- SRR10716763 for female genital discs, 48h after puparium formation

- SRR10716764 for male genital discs, 24h after puparium formation

- SRR10716765 for female genital discs, 24h after puparium formation

- SRR10716766 for adult female tarsae

- SRR10716767 for adult female proboscis + maxillary palps

- SRR10716768 for adult female ovipositor

- SRR10716770 for adult female antennae

- SRR10716771 for adult male antennae

### Individual Whole Genome Shot-Gun data (SRA accessions numbers)

#### New to this study

- SRR10260311 for the female individual mtp_f19

- SRR10260312 for the male individual mtp_m19

### Pool Whole Genome Shot-Gun data (SRA accessions numbers)

#### New to this study

- SRR10260310 for the WT3-2.0 pool

#### From Olazcuaga et al. (in prep.)

- SRR10260026 for the US-Wat pool

- SRR10260031 for the US-Haw pool

- SRR10260027 for the CN-Nin pool

#### From Chiu et al. (2013)

- SRR942805 for the WT3-1.0 pool

## AUTHOR CONTRIBUTIONS

AE, MG, MP, NG and BP conceived the project, MP, NG, AE, MG and BP designed the experiments. RB assembled the genome, MP did the genome alignments, MP and RB annotated and analyzed the genome assembly. HP prepared the samples for Pool-seq, MG analyzed the Pool-seq data. JG and MK prepared the samples used for RNA-seq. MP, AE, MG and BP wrote the manuscript. AE, BP, MG, NG secured funding.

