## Supplementary figures. for "Near-chromosome level genome assembly of the fruit pest *Drosophila suzukii* using long-read sequencing"

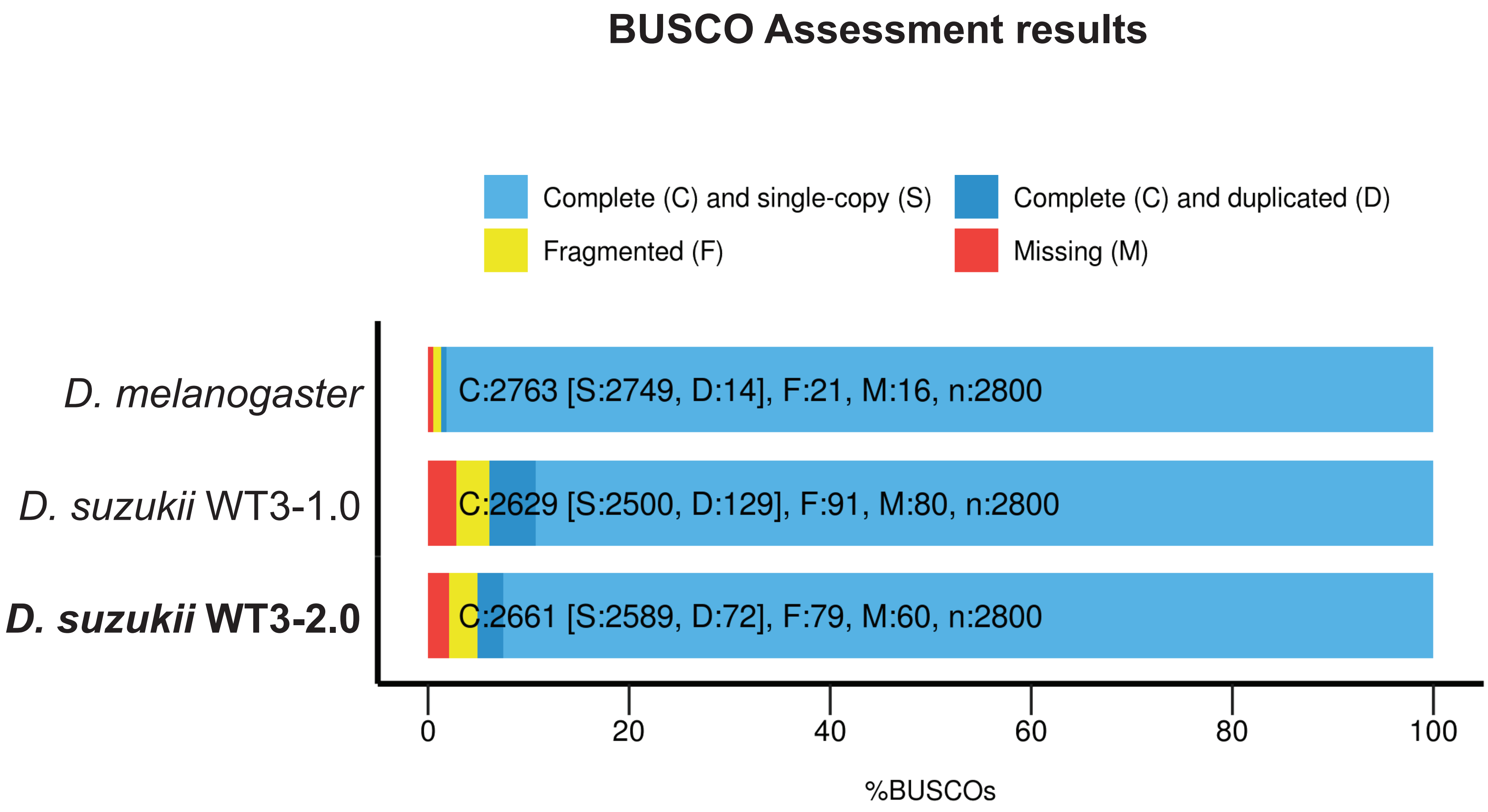
BUSCO Assessment results
Complete (C) and single-copy (S)
Complete (C) and duplicated (D)
Fragmented (F)
Missing (M)
D. melanogaster
D. suzukii WT3-1.0
D. suzukii WT3-2.0
C:2763 [S:2749, D:14], F:21, M:16, n:2800
C:2629 [S:2500, D:129], F:91, M:80, n:2800
C:2661 [S:2589, D:72], F:79, M:60, n:2800
0
20
40
60
80
100
%BUSCOs

FIGURE S1

Figure S1: BUSCO statistics for the different assembly steps.

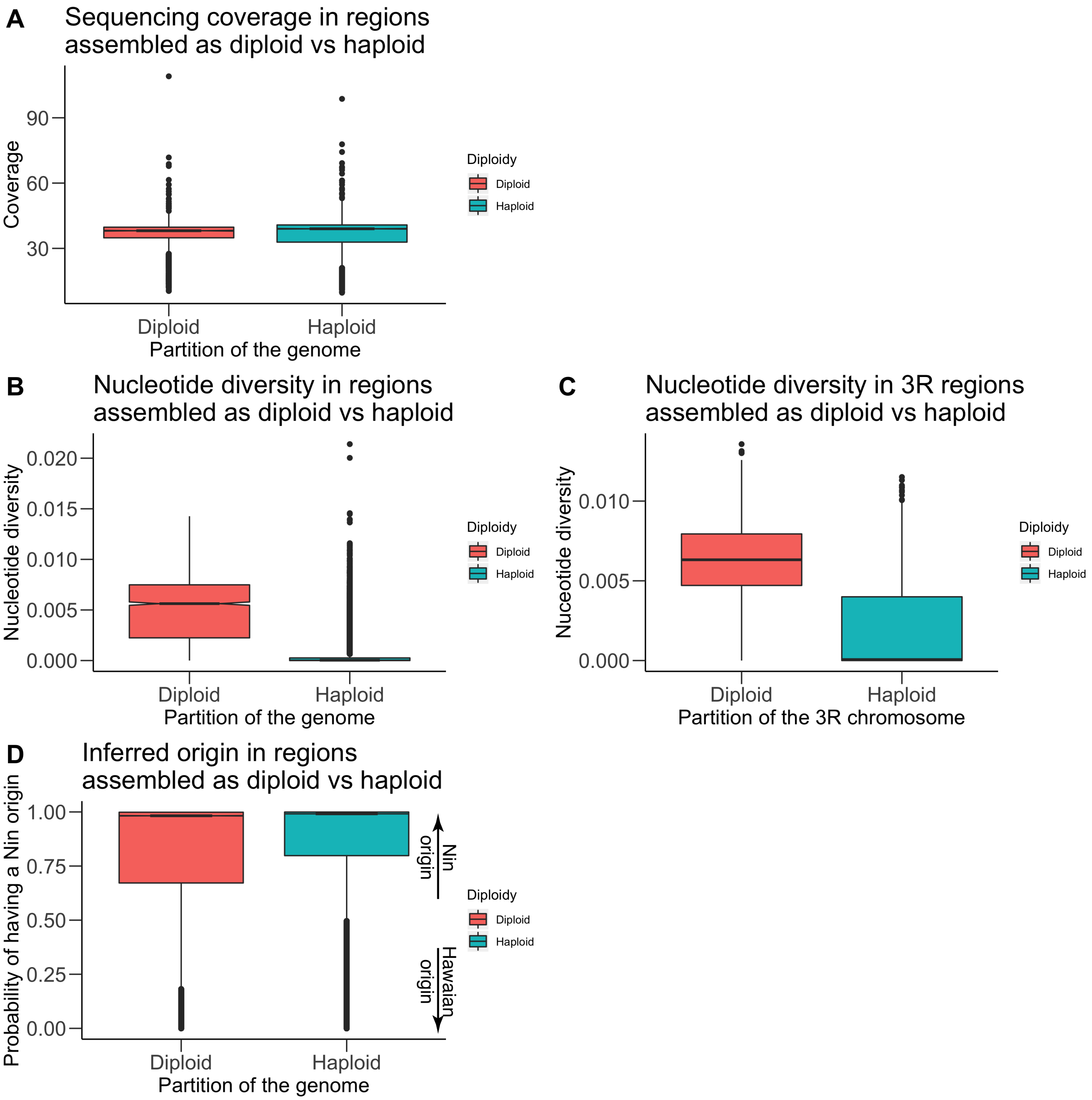
A
Sequencing coverage in regions
assembled as diploid vs haploid
Coverage
90
60
30
Diploid
Haploid
Partition of the genome
Diploidy
Diploid
Haploid
B
Nucleotide diversity in regions
assembled as diploid vs haploid
Nucleotide diversity
0.020
0.015
0.010
0.005
0.000
Diploid
Haploid
Partition of the genome
Diploidy
Diploid
Haploid
C
Nucleotide diversity in 3R regions
assembled as diploid vs haploid
Nuceotide diversity
0.010
0.005
0.000
Diploid
Haploid
Partition of the 3R chromosome
Diploidy
Diploid
Haploid
D
Inferred origin in regions
assembled as diploid vs haploid
Probability of having a Nin origin
1.00
0.75
0.50
0.25
0.00
Nin origin
Hawaiian origin
Diploid
Haploid
Partition of the genome
Diploidy
Diploid
Haploid

FIGURE S2

Figure S2 : Comparison of coverage (**A**) and nucleotide diversity (**B**) between regions for which an alternative sequence was assembled (« Diploid ») and the rest of the genome (« Haploid »). Reads from the WT3-2.0 Pool-seq sample were used and mean coverage was calculated on 10 kb windows. (**C**) Same analysis than in (**B**) for the contigs associated with chromosomal arm 3 only. (**D**) Probability of being of population CN-NIN (Ningbo, China) origin for SNPs located within either the diploid or the haploid regions of the genome. A value close to 1 means that the SNP is predicted as being in a genomic region of CN-NIN origin and a value close to 0 means that the SNP is predicted as being in a region of alternative populational origin (*i.e.*, US-Haw for Hawaii – USA).

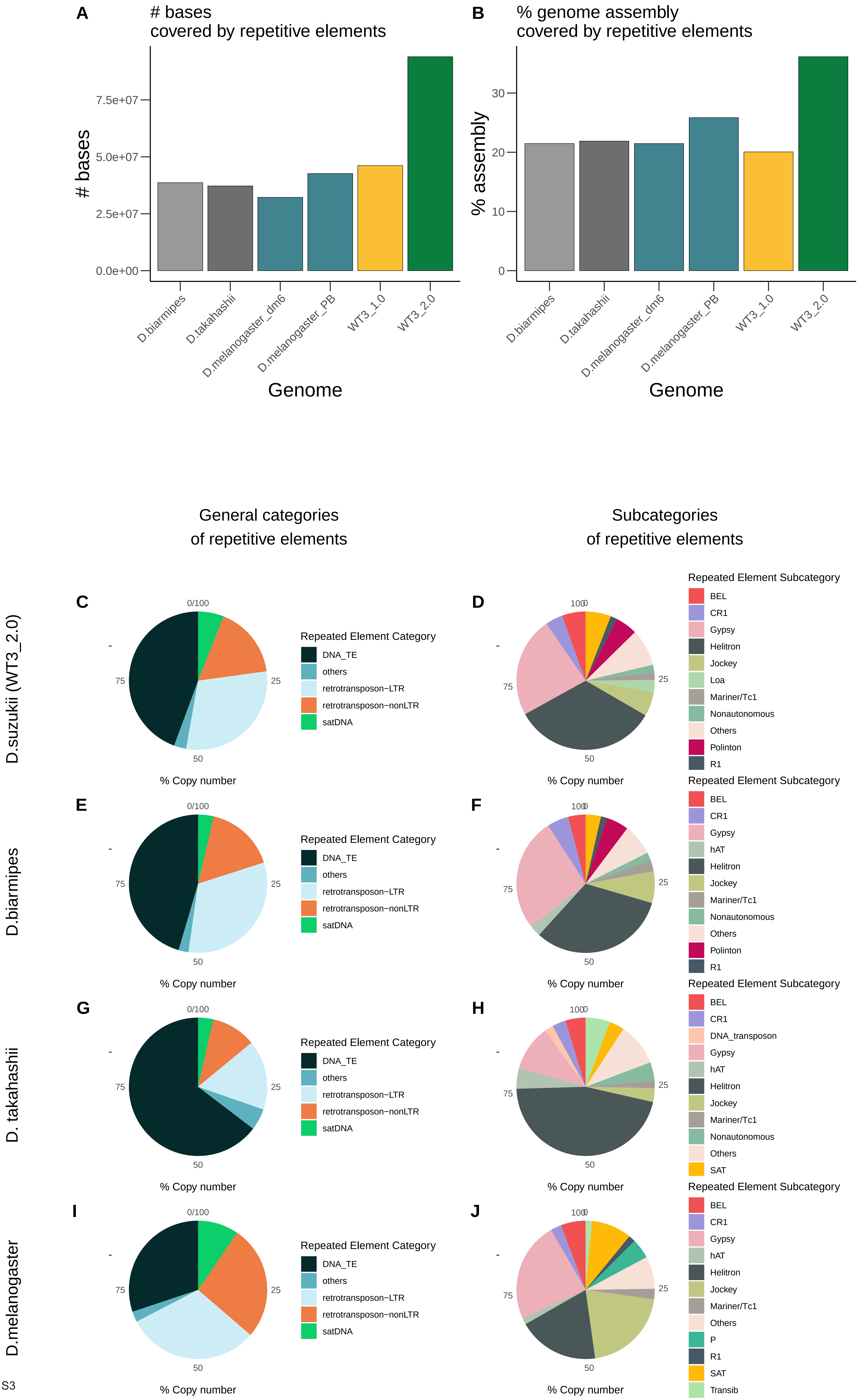
A
### bases
covered by repetitive elements
### bases
7.5e+07
5.0e+07
2.5e+07
0.0e+00
D.biarmipes
D.takahashii
D.melanogaster_dm6
D.melanogaster_PB
WT3_1.0
WT3_2.0
Genome
B
% genome assembly
covered by repetitive elements
% assembly
30
20
10
0
D.biarmipes
D.takahashii
D.melanogaster_dm6
D.melanogaster_PB
WT3_1.0
WT3_2.0
Genome
General categories
of repetitive elements
Subcategories
of repetitive elements
D.suzukii (WT3_2.0)
C
0/100
25
50
75
% Copy number
Repeated Element Category
DNA_TE
others
retrotransposon−LTR
retrotransposon−nonLTR
satDNA
D
100
25
50
75
% Copy number
Repeated Element Subcategory
BEL
CR1
Gypsy
Helitron
Jockey
Loa
Mariner/Tc1
Nonautonomous
Others
Polinton
R1
D.biarmipes
E
0/100
25
50
75
% Copy number
Repeated Element Category
DNA_TE
others
retrotransposon−LTR
retrotransposon−nonLTR
satDNA
F
100
25
50
75
% Copy number
Repeated Element Subcategory
BEL
CR1
Gypsy
hAT
Helitron
Jockey
Mariner/Tc1
Nonautonomous
Others
Polinton
R1
D. takahashii
G
0/100
25
50
75
% Copy number
Repeated Element Category
DNA_TE
others
retrotransposon−LTR
retrotransposon−nonLTR
satDNA
H
100
25
50
75
% Copy number
Repeated Element Subcategory
BEL
CR1
DNA_transposon
Gypsy
hAT
Helitron
Jockey
Mariner/Tc1
Nonautonomous
Others
SAT
D.melanogaster
I
0/100
25
50
75
% Copy number
Repeated Element Category
DNA_TE
others
retrotransposon−LTR
retrotransposon−nonLTR
satDNA
J
100
25
50
75
% Copy number
Repeated Element Subcategory
BEL
CR1
Gypsy
hAT
Helitron
Jockey
Mariner/Tc1
Others
P
R1
SAT
Transib

FIGURE S3

Figure S3: Comparison of the repetitive sequences repertoire between *D suzukii*, *D, melanogaster*, *D. biarmipes* and *D. takahashii.* Comparison of (**A**) the number of bases and (**B**) the percentage of the genome assembly covered by repetitive elements. Two assemblies were used for *D. suzukii* (WT3-1.0 and WT3-2.0) and for *D. melanogaster* (dm6 and an assembly done using PacBio (PB) reads), (**C-J**) The number of copies of each type of repetitive elements was compared for broad categories (**C**, **E**, **G** and **I**) and more specific categories (**D, F**, **H** and **J**) of repetitive elements.

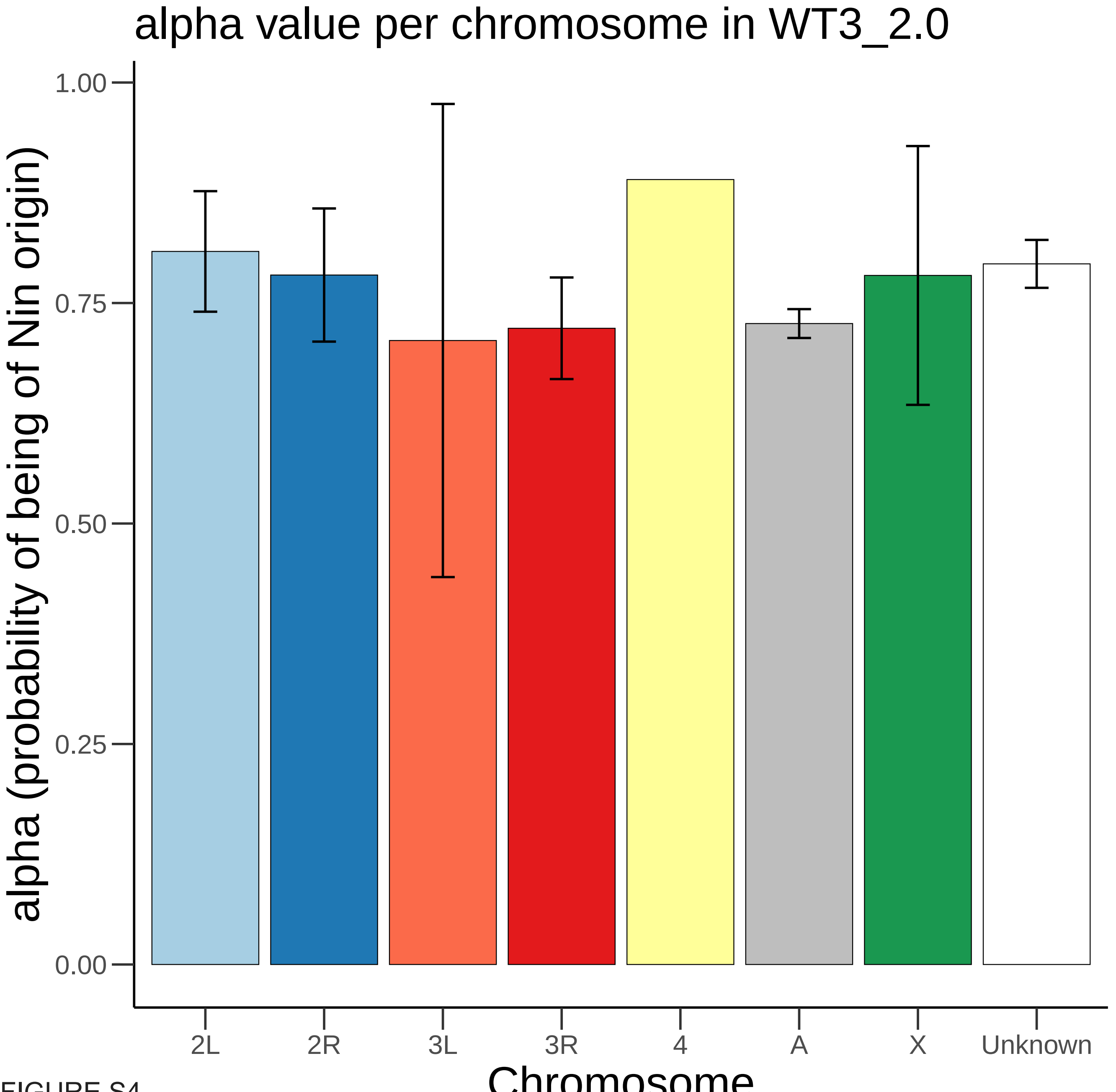
alpha value per chromosome in WT3_2.0
alpha (probability of being of Nin origin)
1.00
0.75
0.50
0.25
0.00
2L
2R
3L
3R
4
A
X
Unknown
Chromosome

FIGURE S4

Figure S4: Chromosomal distribution of alpha, the probability of being of population CN-NIN (Ningbo, China). Values correspond to the relative length proportion of each contig with a Nin origin. “2L”, “2R”, “3L”, “3R”, “4”, “X”: assigned *D. melanogaster* chromosome for each WT3-2.0 contigs. “A”: autosomal contigs with no clear corresponding *D. melanogaster* chromosome. “Unknown”: contigs for which chromosomal features (i.e., assignment to autosomal or X chromosomes and to *D. melanogaster* chromosome arm) remain unknown. Error bars correspond to S.E.M.

Table S1: Descriptive statistics of the *D. suzukii* assemblies WT3-2.0 and WT3-1.0, and of the *D. melanogaster* assembly dm6.

Table S2: Gene orthology table based on genome alignment between WT3-2.0, WT3-1.0 (Chiu et al., 2013) and *D. melanogaster* r6.03.
