## Supplementary file 2 for "Near-chromosome level genome assembly of the fruit pest *Drosophila suzukii* using long-read sequencing"

### Supplementary Text S2

#### *The Hidden Markov Model for assembly painting*

We here describe the one-order Hidden Markov Model (HMM) we developed to model the *Drosophila suzukii* assembly WT3-2.0 as a mosaic of chromosomal segments from either Chinese ("C") or Hawaiian ("H") origins.

Let $s_i$ represent the (local) ancestral origin ($s_i = C$ or $s_i = H$) of the allele in the reference assembly at SNP position $i$ and $d_i$ the physical distance (in bp) between SNP position $i+1$ and $i$. The four different transition probabilities between the (hidden) ancestral origin $s_i$ and $s_{i+1}$ of two adjacent SNPs then read:

$$\begin{cases} \mathbb{P}\left[s_{i+1} = \mathrm{C} \mid s_i = \mathrm{C}\right] & = e^{-\rho d_i} + \left(1 - e^{-\rho d_i}\right)\alpha \\ \mathbb{P}\left[s_{i+1} = \mathrm{H} \mid s_i = \mathrm{C}\right] & = \left(1 - e^{-\rho d_i}\right)(1-\alpha) \\ \mathbb{P}\left[s_{i+1} = \mathrm{C} \mid s_i = \mathrm{H}\right] & = \left(1 - e^{-\rho d_i}\right)\alpha \\ \mathbb{P}\left[s_{i+1} = \mathrm{H} \mid s_i = \mathrm{H}\right] & = e^{-\rho d_i} + \left(1 - e^{-\rho d_i}\right)(1-\alpha) \end{cases} \tag{1}$$

The parameter $\rho$ corresponds to the per bp ancestry switching rate along the sequence assembly and the parameter $\alpha$ to the proportion of the genome assembly of Chinese ("C") origin. Hence, the term $\left(1 - e^{-\rho d_i}\right)$ is the probability that a change of ancestry occurred between SNP position $i$ and $i+1$, and in that case the reference allele at position $i+1$ is of Chinese (respectively Hawaiian) origin with probability $\alpha$ (respectively $1-\alpha$). It might be tempting to interpret $\rho$ as a historical recombination rate with $\rho = \rho_0(t_a + t_i)$ where $\rho_0$ is the recombination rate per bp and generation; $t_a$ is the time (in generations) separating the WT3 founding population (Watsonville, USA) from the admixture between Chinese and Hawaiian populations; and $t_i$ is the number of generations of inbreeding to create the sequenced WT3 strain. However, because inbreeding in the isofemale strains remains incomplete in several genomic regions, the sequence assembly cannot strictly be viewed as a single haplotype and may rather correspond locally to a consensus of different chromosomal segments (i.e., *in silico* recombinant). As a result, the estimated $\rho$ is expected to provide an upwardly biased estimate of the historical recombination rate. It should also be noticed that the overall length of tracts of alleles from Chinese (resp. Hawaiian) ancestry has an expected mean equal to $\frac{1}{(1-\alpha)\rho}$ (resp. $\frac{1}{\alpha\rho}$) as consecutive segments in the sequence assembly might belong to the same ancestry.

To complete the specification of the HMM, the emission probability for the observed reference allele $a_i$ at SNP position $i$ given the ancestral origin $s_i = j$ was set equal to its frequency in the corresponding ancestry $j$ (i.e., $j = C$ or $j = H$) as estimated from the

Pool-Seq data, i.e. $\mathbb{P}[a_i \mid s_i = j] = \widehat{f_{ij}}$. The estimated frequencies $\widehat{f_{ij}}$ of the reference allele (i.e., the one of the assembly) were computed using a Laplace estimator as: $\widehat{f_{ij}} = \frac{r_{ij}+\epsilon}{c_{ij}+\epsilon}$ where $r_{ij}$ and $c_{ij}$ represent the observed read count for the reference allele and overall coverage for SNP $i$ in population $j$. The parameter $\epsilon$ controls the prior Beta distribution for allele frequency (namely $\beta(\epsilon, \epsilon)$ which is the expected distribution at the mutation-drift equilibrium for a constant size population with $\theta = 4N_e\mu = \epsilon$) and was set to 0.01. Note that the main rationale for using such an estimator was to avoid fixed allele frequencies estimates, i.e., equal to 0 or 1 (the minimal possible minor allele frequency being equal to 0.005) which would lead to absorbing states in the painting model. In other words, an allele unobserved in the Pool-Seq reads was considered as extremely rare rather than completely absent from the corresponding population.

Given a set of parameter values, we used the forward-backward algorithm (Rabiner, 1989) to compute the model likelihood (i.e., the conditional probability of the sequence of reference alleles at each and every SNP positions) and the probability to belong to the two ancestries at each SNP position by integrating over all possible sequences of ancestries. Note that the contigs were randomly ordered and the distances between markers belonging to different contigs was set to an arbitrary large value equal to $10^{10}$. To estimate the parameter values that maximized the likelihood, we then used the default Nelder and Mead method as implemented in the optim function from the R stats package (R Core Team, 2013). In order to work with unconstrained parameters in the optimization procedure, the model was reparameterized with the following two new parameters, $\rho^\star = \log(\rho)$ and $\alpha^\star = \log(1-\alpha) - \log(\alpha)$. Standard errors of the resulting Maximum Likelihood Estimates (MLEs) for the two HMM parameters were obtained from the approximate Hessian of minus the log-likelihood (Zucchini & MacDonald, 2009, , p.53).

We further computed estimates for each contig $k$, with a number of analyzed SNPs $n_k > 2$, both the estimated proportion of Chinese origin $\widehat{\alpha_k}$ and the ancestry change rate $\widehat{\rho_k}$ based on the (posterior) expected number of chunks from Chinese ($\widehat{\eta_k^{(C)}}$) or Hawaiian ($\widehat{\eta_k^{(H)}}$) origin and their corresponding overall expected length ($\widehat{\lambda_k^{(C)}}$ and $\widehat{\lambda_k^{(H)}}$ respectively) as:

$$\widehat{\alpha_k} = \frac{\widehat{\lambda_k^{(C)}}}{\widehat{\lambda_k^{(C)}} + \widehat{\lambda_k^{(H)}}} \quad \text{and} \quad \widehat{\rho_k} = \frac{\widehat{\eta_k^{(C)}} + \widehat{\eta_k^{(H)}}}{\widehat{\lambda_k^{(C)}} + \widehat{\lambda_k^{(H)}}} \tag{2}$$

More precisely, following Lawson *et al* (2012) (eqs. 3 and 4 in Text S1), $\widehat{\lambda_k^{(C)}}$ and $\widehat{\lambda_k^{(H)}}$ were computed as:

$$\widehat{\lambda_k^{(C)}} = \frac{1}{2}\sum_{j=2}^{n_k}\left(\widehat{p_{k,j-1}} + \widehat{p_{k,j}}\right)d_{j-1} \quad \text{and} \quad \widehat{\lambda_k^{(H)}} = \frac{1}{2}\sum_{j=2}^{n_k}\left(2 - \widehat{p_{k,j-1}} + \widehat{p_{k,j}}\right)d_{j-1} \tag{3}$$

and $\widehat{\eta_k^{(C)}}$ and $\widehat{\eta_k^{(H)}}$ were computed as:

$$\widehat{\eta_k^{(C)}} = \widehat{p_{k,1}} + \sum_{j=2}^{n_k} \widehat{\tau_j^{(C)}} \qquad \text{and} \qquad \widehat{\eta_k^{(H)}} = 1 - \widehat{p_{k,1}} + \sum_{j=2}^{n_k} \widehat{\tau_j^{(H)}} \tag{4}$$

where, for the $j$th SNP position of contig $k$, $\widehat{p_{k,j}}$ is the posterior probability that the reference allele on the assembly is of Chinese origin, and $\widehat{\tau_j^{(C)}}$ (respectively $\widehat{\tau_j^{(H)}}$) represents the posterior probability that the reference assembly allele is of Chinese (resp. Hawaiian) origin given at least one change of ancestry occurred between SNP position $j-1$ and $j$. The $\widehat{p_{k,j}}$'s, $\widehat{\tau_j^{(C)}}$'s and $\widehat{\tau_j^{(H)}}$'s posterior probabilites were all estimated using the forward-backward algorithm (Rabiner, 1989) setting the genome-wide parameters $\rho$ and $\alpha$ to their MLE values.
