## Supplementary material for "Near-chromosome level genome assembly of the fruit pest *Drosophila suzukii* using long-read sequencing": Table S1

| | *D. suzukii* WT3-2.0 | *D. suzukii* WT3-1.0 | *D. melanogaster dm6* |
|---|---|---|---|
| **# contigs** | 546 | 4813 | 1870 |
| **Largest contig** | 25 589 241 | 22 559 587 | 32 079 331 |
| **Total length** | 268 012 156 | 231 788 855 | 143 726 002 |
| **N50** | 2 609 782 | 397 157 | 25 286 936 |
| **L50** | 15 | 73 | 3 |
